# A Distribution-aware Semi-Supervised Pipeline for Cost-effective Neuron Segmentation in Volume Electron Microscopy

**DOI:** 10.1101/2024.05.26.595303

**Authors:** Yanchao Zhang, Hao Zhai, Jinyue Guo, Jing Liu, Qiwei Xie, Hua Han

**Author notes:** Corresponding author (J. Liu); (H. Han). Equal contribution (Yanchao Zhang, Hao Zhai).

## Abstract

Semi-supervised learning offers a cost-effective approach for neuron segmentation in electron microscopy (EM) volumes. This technique leverages extensive unlabeled data to regularize supervised training for more robust predictions of neuron boundaries. However, the distribution mismatch between labeled and unlabeled data, due to limited annotations and diverse neuronal structures, can cause unreliable predictions on unlabeled data, thus limiting the generalization of semi-supervised models. In this paper, we develop a distribution-aware pipeline to address the inherent mismatch issue and enhance semi-supervised neuron segmentation in EM volumes. At the data level, we propose an unsupervised heuristic to select valuable sub-volumes as labeled data based on distribution similarity in a pretrained feature space, ensuring a representative coverage of neuronal structures. At the model level, we introduce an axial-through mixing strategy into anisotropic neuron segmentation and integrate it into a semi-supervised framework. Building on this, we establish cross-view consistency constraints through intra- and inter-mixing of labeled and unlabeled data, which facilitates the learning of shared semantics across distributions. Extensive experiments on public EM datasets from multiple species, imaging modalities, and resolutions demonstrate the effectiveness of our method. Codes and demos are available at https://github.com/yanchaoz/SL-SSNS.

## 1. Introduction

Connectomics aims to uncover brain mechanisms and intelligence by reconstructing and interpreting neural circuits at the synaptic level (Ngai, 2022). Currently, only volume electron microscopy (EM), with its nanoscale spatial resolution, supports imaging the intricate morphology of neurons and individual synapses in brain tissue (Peddie et al., 2022; Peddie and Collinson, 2014). However, this technique generates massive volumetric data ranging from several hundred terabytes to petabytes (Abbott et al., 2020; Shapson-Coe et al., 2024; Zheng et al., 2018; Dorkenwald et al., 2023; Scheffer et al., 2020), which calls for automated methods to accelerate data processing and analysis. Therefore, efficient and accurate neuron reconstruction from EM volumes has become critical to advance the digitalization of the complete atlas of the nervous system.

Recent advances in learning-based neuron segmentation have shown promising results across various EM modalities and voxel resolutions (Lee et al., 2017; Januszewski et al., 2018; Lee et al., 2021; Funke et al., 2018; Sheridan et al., 2022). Typically, these methods rely on convolutional neural networks to predict neuronal membrane descriptors, followed by supervoxel extraction and agglomeration to obtain final segmentation. Despite their success, a fundamental challenge persists: The pronounced heterogeneity of neuronal structures across EM volumes necessitates broad distributional coverage in labeled data for robust generalization, yet the prohibitive cost of expert annotation severely limits its availability. For example, the fly hemi-brain segmentation model was trained on merely 0.003% of the entire dataset (0.8GB out of 26TB) (Scheffer et al., 2020). This inherent tension between data diversity and annotation scarcity poses a major barrier to effective model training.

Semi-supervised learning has emerged as a promising paradigm to mitigate annotation scarcity by leveraging abundant unlabeled data to enhance model performance. Related works in biomedical image segmentation have developed techniques, such as consistency learning (Huang et al., 2022; Luo et al., 2021; Yu et al., 2019) and self-training (Wang et al., 2022b; Shi et al., 2021), to generate supervisory signals from unlabeled data. Notably, these methods typically assume that labeled and unlabeled data are drawn from the same distribution, thereby allowing unlabeled data to effectively guide network optimization. In practice, this assumption often fails, particularly in dense neuron segmentation, due to the complexity of neural structures, large-scale unlabeled volumes, and limited annotation coverage, as illustrated in Figure 1. Distribution mismatches can cause unreliable predictions on unlabeled data, resulting in suboptimal semi-supervised learning and degraded segmentation performance.

**Figure 1.**
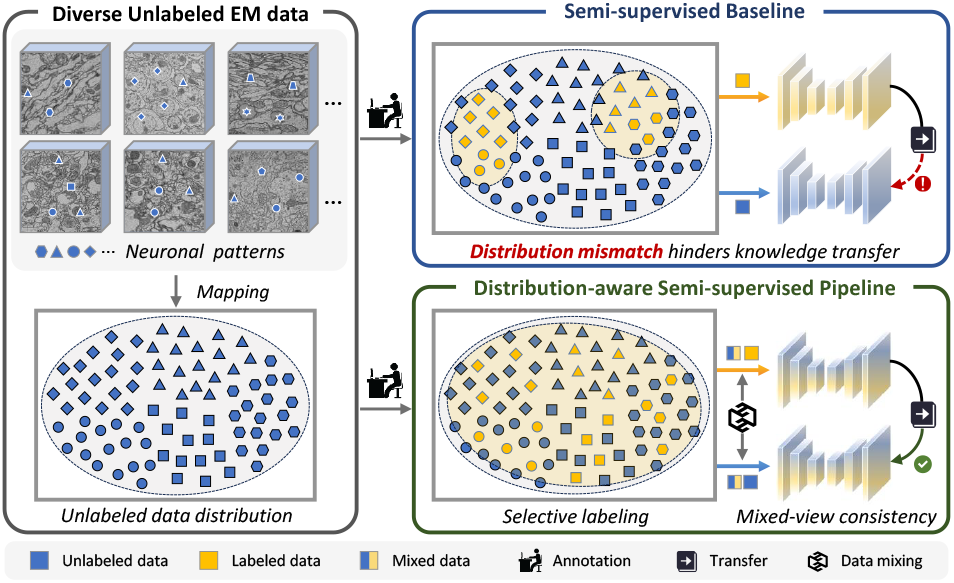
Limited by annotation budgets, labeled data often fails to capture the distribution of the unlabeled set. This distribution mismatch impedes knowledge transfer and degrades segmentation performance. Our proposed approach mitigates this issue by aligning the labeled and unlabeled distributions through coverage-based selective labeling and mixed-view consistency learning.

Addressing the distribution mismatch issue requires joint consideration at both the data and model levels. **At the data level**, an intuitive yet underexplored direction involves selecting sub-volumes that effectively capture the distributional diversity of the entire EM volume. Two main challenges arise in this context: (1) Lack of supervision — In a fully unsupervised setting, the absence of any initial annotations complicates the identification of informative regions worth labeling (Zheng et al., 2019; Jin et al., 2022). (2) Continuous data structure — Unlike discrete-instance datasets (e.g., ImageNet (Russakovsky et al., 2015)) where diversity sampling is straightforward (Wang et al., 2022a; Mannix and Bondell, 2023), EM volumes are continuous in space, making it nontrivial to evaluate the representativeness of sub-regions with varying sizes. **At the model level**, narrowing the distribution gap requires mechanisms that promote bidirectional information flow between labeled and unlabeled data. However, many existing approaches treat labeled and unlabeled samples independently, resulting in asymmetric feature learning and limited generalization. Building effective connections between distributions remains a central challenge in semi-supervised neuron segmentation.

In this work, we propose a distribution-aware pipeline for more effective semi-supervised neuron segmentation, as shown in Figure 1. Specifically, we first perform self-supervised learning on the unlabeled EM dataset to develop an embedding network that provides semantically meaningful representations for local patches. Building on this, we propose a quantitative heuristic for selecting representative sub-volumes by aggregating spatially adjacent patches under a coverage-driven criterion in the embedding space. This strategy enables the identification of sub-volumes that faithfully capture the underlying data distribution. Afterward, we train a segmentation model using selectively labeled sub-volumes and the unlabeled data in a semi-supervised manner. By adapting the spatial mixing strategy to anisotropic EM data, we generate transitional inputs that bridge the feature space between distributions, facilitating shared semantic learning through mixed-view consistency regularization. This two-stage approach effectively reduces the distribution gap both in the input and feature spaces, demonstrating practical value in large-scale neuron reconstruction. Our contributions are summarized as follows:

1. We propose a unified, distribution-aware semi-supervised pipeline that mitigates mismatches between labeled and unlabeled distributions at both data selection and model training stages.
2. We develop an unsupervised, quantitative heuristic for selecting representative sub-volumes from EM data, ensuring comprehensive coverage of structural diversity under limited annotation.
3. We make the first attempt to integrate spatial mixing into semi-supervised neuron segmentation, which establishes a connection between labeled and unlabeled datasets and enables mixed-view consistency regularization.
4. We validate our approach on diverse EM datasets across various species and resolutions, demonstrating strong generalization and providing new insights into cost-effective neuron segmentation.

## 2. Related work

### 2.1 Neuron Segmentation

As an instance segmentation problem, neuron reconstruction aims to assign a unique label to every voxel in the EM volumes that belongs to the same neuron. This task is challenging due to the unprecedented scale of EM data, the varied textures of neuronal structures, and the scarcity of expert annotations. Advanced methods can be roughly categorized into two types: boundary-based and object-based. Specifically, boundary-based approaches first use 3D U-Net (Çiçek et al., 2016) variants to predict descriptors for neuron boundaries (e.g., voxel affinity graphs, dense voxel embeddings (Lee et al., 2021), and local shape descriptors (Sheridan et al., 2022)). After over-segmentation via watershed transform, graph-based agglomeration is adopted to group supervoxels for instance results (Beier et al., 2017; Funke et al., 2018; Li et al., 2024) (Figure 2(d)). In contrast, object-based approaches, as seen in (Januszewski et al., 2018; Meirovitch et al., 2019), extend the segmented area from the seed points to complete neurites iteratively. While effective, large-scale neuron reconstructions still demand years of manual proofreading before they can support reliable connectivity analysis (Zheng et al., 2018). Enhancing the accuracy of automatic segmentation is therefore crucial for cost-effective connectomics. In this study, we adopt a boundary-based pipeline that employs a 3D CNN to predict voxel affinities along the *x, y*, and *z* axes. To improve affinity predictions, we leverage both limited labeled data and abundant unlabeled data within a semi-supervised framework, aiming to boost model generalization and reduce the burden of manual proofreading.

**Figure 2.**
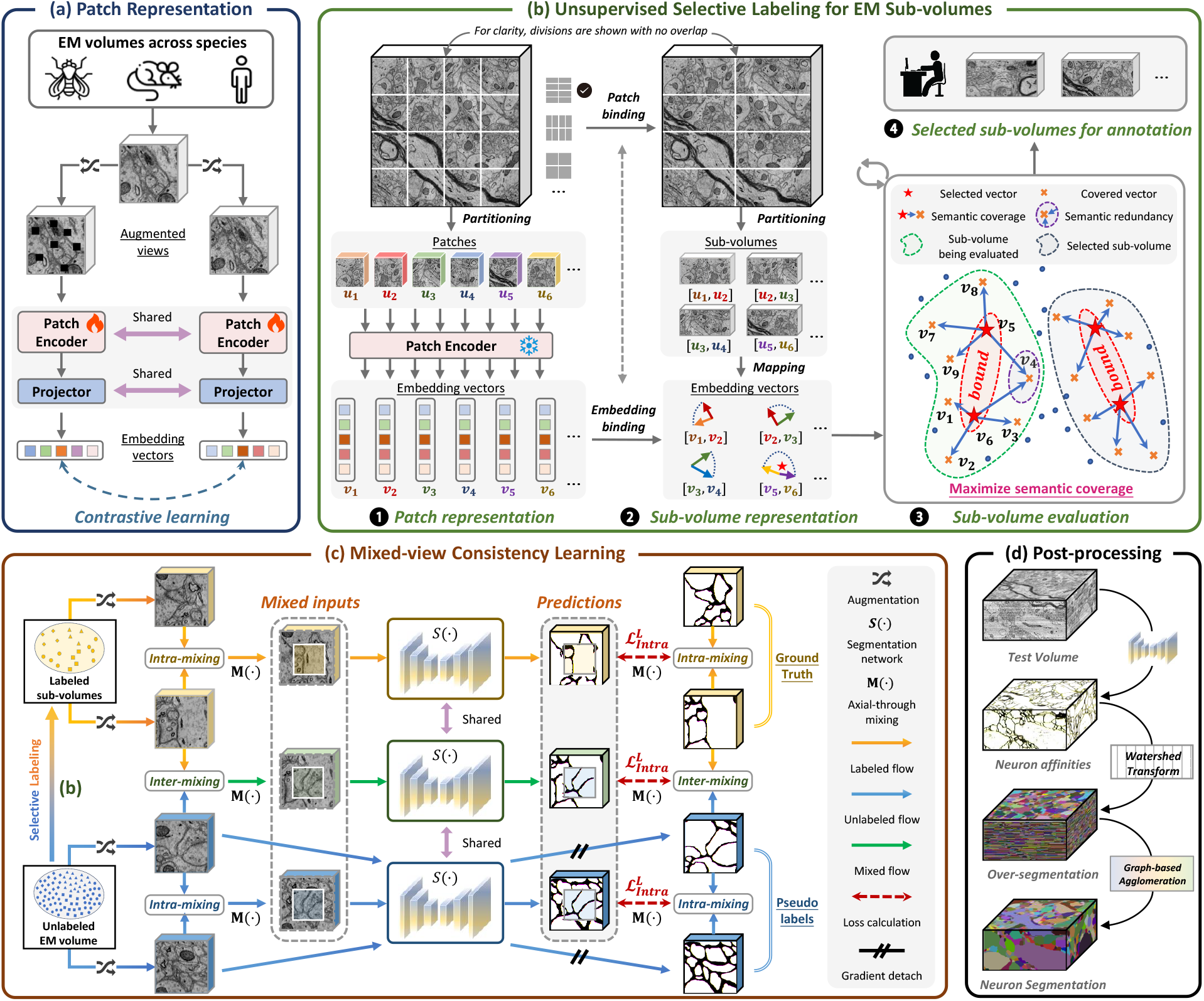
Pipeline overview. (a) A 3D EM patch encoder is trained via contrastive learning across multiple data distributions. (b) After partitioning, the patch encoder maps unlabeled data to the embedding space for EM representation. Then, a quantitative heuristic is designed to identify representative sub-volumes as labeled data iteratively. (c) These sub-volumes are fully exploited via intra-and inter-dataset mixing, with mixed-view consistency enforced to promote shared semantic learning. (d) The trained model first predicts neuron affinities, followed by watershed over-segmentation and graph-based agglomeration for final segmentation.

### 2.2 Semi-supervised Segmentation

Labeled data typically represent only a small fraction of the total imaging volume in practical reconstruction tasks. Semi-supervised learning (SSL) offers a promising solution to this annotation bottleneck by leveraging unlabeled data to enhance model generalization. Self-training and consistency regularization represent two of the most prominent strategies in SSL for generating supervision signals from unlabeled data. Self-training assigns pseudo-labels to unlabeled samples, which are then combined with labeled data for iterative model retraining (Wang et al., 2022b; Shi et al., 2021; Yang et al., 2022). Conversely, consistency regularization is based on the premise that model predictions for unlabeled data should remain stable under various perturbations (Tarvainen and Valpola, 2017; Yu et al., 2019; Luo et al., 2021; Wu et al., 2021). Notably, SSNS-Net (Huang et al., 2022) represents a pioneering application of consistency learning for label-efficient neuron segmentation, demonstrating the potential of semi-supervised approaches.

SSL typically assumes that labeled and unlabeled data share the same underlying distribution (Kurakin et al., 2019; Berthelot et al., 2019). However, this assumption often fails in neuron segmentation due to the intrinsic variability of neural structures and limited annotation budgets. Despite its critical impact, distribution mismatch remains largely underexplored in this domain. This paper proposes a distribution-aware semi-supervised pipeline that addresses domain mismatch through representative data selection and mixed-view consistency learning. Together, these strategies reduce distribution divergence and significantly improve semi-supervised neuron segmentation.

### 2.3 Selective Labeling

Selective labeling (SL), a core focus of active learning, aims to annotate a valuable subset of an unlabeled dataset to optimize supervised learning under limited annotation budgets. Active learning methods are generally categorized into iterative active learning (ITAL) and one-shot active learning (OSAL) based on whether samples are selected iteratively or in a single batch. ITAL has been extensively studied and typically involves iterative selection of informative samples based on uncertainty (Beluch et al., 2018; Konyushkova et al., 2019; Wen et al., 2018) or diversity (Yang et al., 2017; Lin et al., 2020; Sener and Savarese, 2018), with continual network fine-tuning as additional labeled data accrue. However, ITAL methods seldom address the critical initial sample selection problem when no trained model exists, and the iterative cycle of training, inference, and annotation incurs high computational costs, especially for large-scale neuron segmentation. In contrast, OSAL selects all valuable samples in one shot before model training. Typically, these methods first estimate the unlabeled data distribution via unsupervised representation learning, projecting raw data into low-dimensional embeddings, followed by customized strategies (Zheng et al., 2019; Jin et al., 2022; Wang et al., 2022a; Kolluru et al., 2021; Lou et al., 2022) to identify representative and informative samples. Despite these advances, selective labeling for volumetric data such as EM volumes remains underexplored.

Due to the spatial continuity and substantial size variability of neuronal structures, practical reconstruction typically utilizes sub-volumes with adaptable dimensions, from which local patches are sampled for training. Existing OSAL approaches, however, operate on discrete instances, limiting annotation flexibility and failing to capture the continuous nature of neuron boundaries, which in turn degrades segmentation performance. To bridge this gap, we propose a novel OSAL method that enables the selection of valuable sub-regions with flexible sizes within continuous unlabeled EM volumes, thereby better representing the overall data distribution under limited annotation budgets.

## 3. Method

As illustrated in Figure 1, we seek to mitigate the distribution mismatch between labeled and unlabeled datasets to improve semi-supervised neuron segmentation. The proposed approach highlights two primary components: selective labeling of valuable sub-volumes (Section 3.1) and semi-supervised training via mixed-view consistency regularization (Section 3.2), trying to tackle the mismatch issue from data and model levels, respectively.

### 3.1 Unsupervised Selective Labeling for EM Sub-volumes

Given an unlabeled dataset *D*_*u*_ and a limited labeling budget, our objective is to identify and annotate representative sub-volumes (*D*_*l*_) that sufficiently reflect the underlying data distribution. To achieve this, we propose a heuristic method that combines patch-level representation learning with a coverage-based selection criterion.

#### 3.1.1 Unsupervised Representation Learning for EM Patches

To begin, we employ self-supervised learning to train an embedding network that produces low-dimensional, semantically rich representations of 3D EM patches. To promote robust generalization across diverse neuronal patterns, the model is trained on electron microscopy datasets spanning three species: fly, rat, and human. Specifically, the framework, based on Siamese networks (Chen and He, 2021), is optimized by maximizing the similarity between two augmented views of the same volumetric image in the learned feature space, as illustrated in Figure 2(a). We carefully apply perturbations that preserve semantic content, including reflections, rotations, Gaussian blurring, Gaussian noise, simulated misalignment, photometric changes, and random pixel masking. During training, each input volume *u* ∈ ℝ^*D*×*H*×*W*^ is sampled from the unlabeled dataset *D*_*u*_ and randomly augmented to produce two views,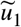and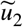. These views are then passed through a Siamese network consisting of two weight-sharing patch encoders, denoted *f* (·), two weight-sharing projectors, denoted as *g*(·), and a predictor, denoted as *h*(·). The network parameters are updated by minimizing the symmetrized loss function:

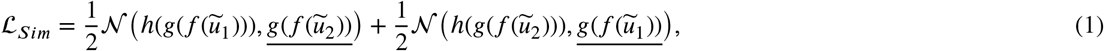

where𝒩 (·, ·) represents the negative cosine similarity between two representations. The underlined terms are treated as constants during backpropagation and do not contribute gradients. The trained patch encoder *f*_*θ*_(·) subsequently serves as an embedding network that projects EM patches into an *L*-dimensional feature space.

### 3.1.2 Coverage-based Sub-volume Selection

This section proposes a coverage-based heuristic to identify representative and informative sub-volumes within the embedding space. For clarity, we distinguish between a patch and a sub-volume: a patch is the smallest unit used for representation learning, whereas a sub-volume is an arbitrarily sized sub-region cropped from the EM volume. The sub-volume annotation strategy is more flexible and suitable for neuron segmentation, addressing the limitations of patch-level annotation, which often fails to capture the spatial continuity and size variability of neuronal structures. Each candidate sub-volume is viewed as a structured aggregation of spatially neighboring patches, and their corresponding embeddings jointly define the representation of the sub-volume in the learned feature space. Carefully selected sub-volumes that cover diverse and typical patches provide a practical approximation of the underlying data distribution.

Formally, the unlabeled volume(s) can be represented as the universal set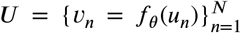, where each *u*_*n*_ ∈ ℝ^*L*^ is a 3D input patch extracted via sliding window partitioning, and *ν*_*n*_ denotes its corresponding embedding vector. Notably, this approach can be readily adapted to datasets of varying scales by adjusting the stride used in patch partitioning, thereby effectively mapping continuous volumetric data into discrete representations. As shown in Figure 2(b), we represent candidate sub-volume(s) as a subset of the universal set *U*, denoted as the selected set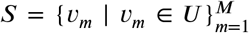, which contains embeddings of spatially contiguous patches extracted from *U*. Intuitively, the distance between the embedding vectors reflects the semantic similarity between their corresponding patches. We thus hypothesize that each selected vector in *S* could represent its *K*-nearest neighbors in *U*, referred to as its covered vectors, thus defining a notion of semantic coverage within the embedding space. Effective sub-volume selection should maximize semantic coverage to represent the data distribution and capture diverse neuronal morphologies. However, semantic redundancy may arise when multiple vectors in *S* cover significantly overlapping embeddings in *U* (see sub-volume evaluation in Figure 2(b)), thereby constraining the overall representational capacity. With this in mind, we introduce the constrained coverage rate (CCR) as a criterion to evaluate the semantic representativeness of the selected set *S*. Formally,

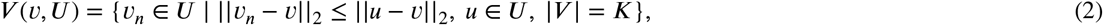

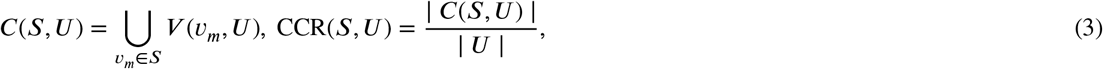

where ||· ||_2_ represent *l*_2_-norm, *V*(*ν, U*) returns the *K*-nearest neighbors of *ν* in *U*, and thus covered set *C* contains all the vectors in *U* that are within the *K*-nearest neighbors of at least one vector in *S*. CCR defines the coverage rate as the percentage of covered vectors in *U*. A higher CCR indicates that the selected sub-volumes cover a larger portion of the neural structures in the entire volume and thus better represent the overall data distribution. Given a labeling budget, our objective is to select a subset *S* ⊆ *U* that maximizes the coverage rate CCR(*S, U*), subject to spatial constraints among the vectors in *S*. Although the optimal subset could theoretically be obtained by exhaustively evaluating all possible sub-volume combinations, this brute-force approach becomes computationally intractable as the dataset size and annotation budget grow. To address this challenge, we propose a coverage-based greedy selection (CGS) algorithm with linear time complexity, offering a practical and scalable solution for large-scale EM volumes. At each iteration, CGS scans the entire volume to generate candidate sub-volumes and extracts their corresponding embedding vectors from *U*. These vectors are provisionally added to the current set *S* to assess its incremental gain in semantic coverage. The sub-volume yielding the largest improvement is then selected for annotation. This process is repeated until the labeling budget is exhausted. Further details are provided in Algorithm 1.

#### Algorithm 1

Coverage-based Greedy Selection for Sub-volumes (CGS)

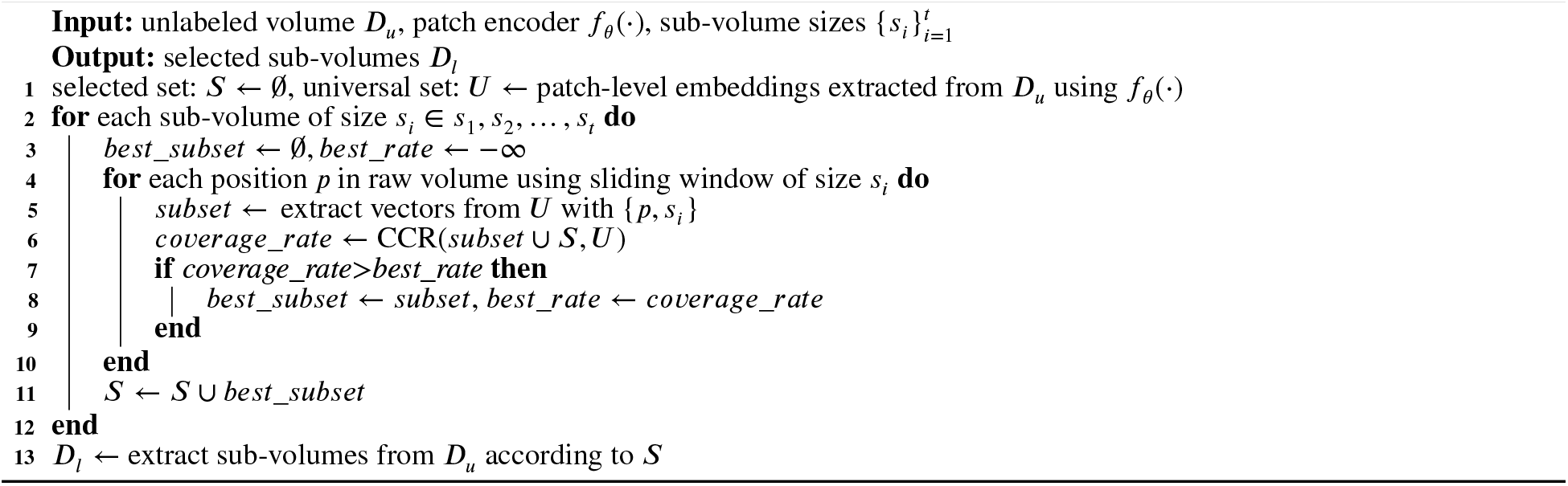

### 3.2 Semi-supervised Neuron Segmentation via Mixed-view Consistency Learning

To mitigate distribution mismatch at the model level, we propose IIC-Net, a semi-supervised framework that promotes feature alignment between labeled and unlabeled data via intra-and inter-dataset mixing (Figure 2(c)).

#### 3.2.1 Axial-through Spatial Mixing for Neuron Segmentation

The affinity graph used to characterize voxel connectivity is generated from dense instance annotations. Each affinity value ranges between 0 and 1, which indicates whether two associated voxels belong to the same instance (1) or not (0) along the *x*/*y*/*z* axis. Let two patches with affinity map be {(*x*_*i*_, *y*_*i*_), (*x*_*j*_, *y*_*j*_)}, where *x*_*i*_, *x*_*j*_ ∈ ℝ^*D*×*H*×*W*^ and *y*_*i*_, *y*_*j*_ ∈ {0, 1}^*C*×*D*×*H*×*W*^. Spatical mixing for raw patches and affinities are denoted as M^*R*^(·) and M^*A*^(·), respectively. Currently, large-scale volume EM data typically exhibits strong anisotropy (Zheng et al., 2018; Shapson-Coe et al., 2024), i.e., the *z*-axis resolution is much lower than that of the *x*-and *y*-axes. To prevent potential semantic ambiguity caused by asymmetric axial and tangential resolution, we employ axial-through masking strategies in spatial mixing, as shown in Figure 3. The mixed result for *x*_*i*_ and *x*_*j*_ is given by

**Figure 3.**
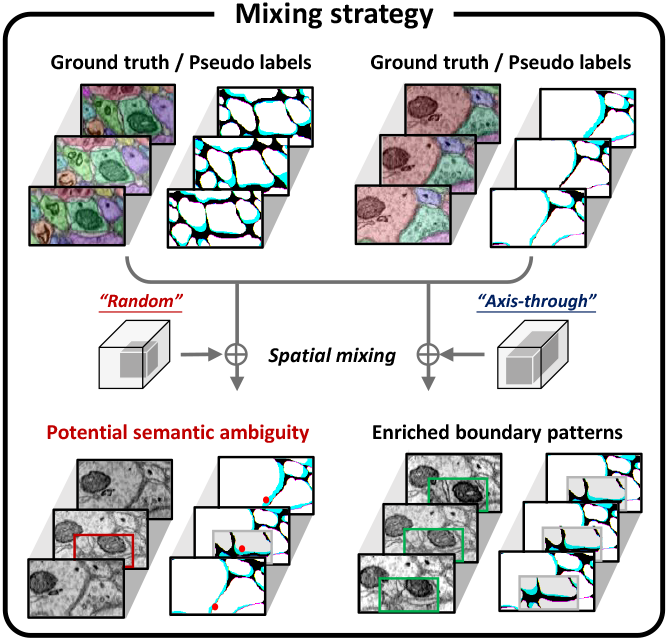
The low axial resolution of anisotropic EM volumes often leads to semantic ambiguity in axial affinities when applying standard random spatial mixing. To alleviate this, we adopt an axial-through mixing strategy, which preserves semantic continuity and avoids abrupt changes in affinity values.

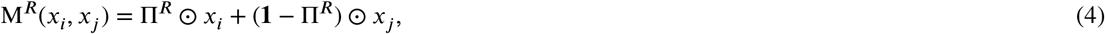

where ⊙ denotes Hadamard product, Π^*R*^ ∈ {0, 1}^*D*×*H*×*W*^ is a binary matrix determined by the mixing strategy, indicating whether the voxel comes from *x*_*i*_ (1) or *x*_*j*_ (0). The size of the zero-value region(s) in Π is *D* × *βH* × *βw*, where *β* ∈ (0, 1). The mixed result for *c*-channel affinities *y*_*i*_ and *y*_*j*_ is computed by

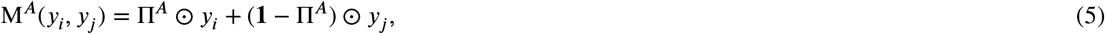

where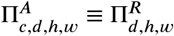, indicating that the same mixing operation is performed on each channel of the affinity graph.

#### 3.2.2 Mixed-view Consistency Learning

Given that network design is not the primary focus of this work, we adopt a modified 3D U-Net as the segmentation backbone *S*(·), which has been widely applied in neuron segmentation (Lee et al., 2017, 2021; Huang et al., 2022). During training, we randomly sample two labeled pairs {(*l*_1_, *y*_1_), (*l*_2_, *y*_2_)} from *D*_*l*_ and two unlabeled volumes {*u*_1_, *u*_2_} from *D*_*u*_. Let M_*i*_ denote the mixing operation associated with a binary mask Π_*i*_. Prior to mixing, we apply random data augmentations to increase the diversity of views, including Gaussian blurring, simulated misalignment, additive Gaussian noise, random masking, and photometric perturbations. As illustrated in Figure 2(c), the supervision signal is built on intra-and inter-dataset mixing. Specifically,

##### Intra-dataset mixing

To maximize the utility of limited annotations and facilitate network adaptation to mixed inputs, the mixing strategy is employed within both labeled and unlabeled datasets. For labeled data, the supervised loss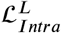is defined as

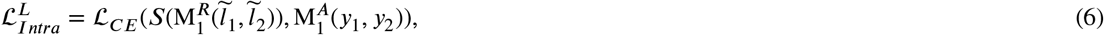

where ℒ_*C E*_ represents cross entropy loss,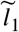and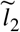are randomly augmented views of *l* and *l*, respectively. We leverage unlabeled data by imposing consistency constraints between predictions of perturbed and original inputs. Specifically, the soft pseudo labels for randomly augmented inputs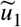and 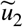 are computed by *S*(*u*_1_) and *S*(*u*_2_), and thus the mixed-view consistency loss for unlabeled data is given by

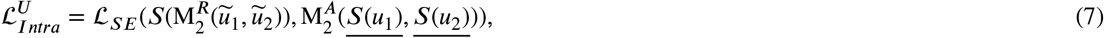

whereℒ_*SE*_ represents square error loss.

##### Inter-dataset mixing

To alleviate asymmetric feature learning caused by distribution gaps between labeled and unlabeled data, we perform spatial mixing across datasets. This generates transitional inputs that enable bidirectional information flow and promote shared semantic learning. In detail, the annotated area in the mixed image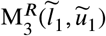is supervised under label map *y*_1_, while the unlabeled area is with the pseudo label *S*(*u*_1_). A binary mask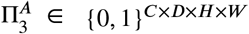and identifies the source of each voxel, i.e., labeled part (1) or unlabeled part (0). Therefore, the supervision signal of the mixed image can be divided into two parts, denoted as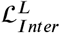 and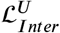. Formally,

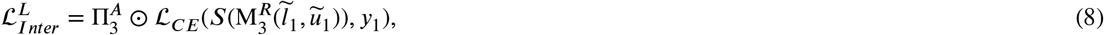

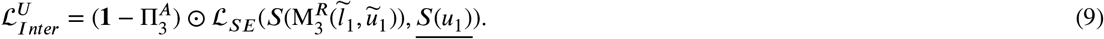

Since expert-annotated masks are generally more accurate than pseudo labels assigned to unlabeled patches, we introduce a weighting coefficient λ to balance their contributions in training. The complete loss function is defined as

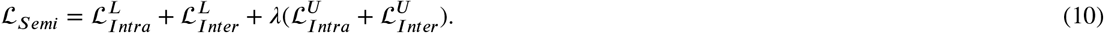

##### Warming-up

Consistency regularization is effective only when applied to confident predictions on unlabeled data. However, achieving this condition can be challenging when the weights of the segmentation network are randomly initialized. To prevent convergence to sub-optimal solutions, we first train the network with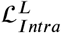, and then fine-tune it with the complete loss function.

## 4. Experiments setup

### 4.1 Dataset

We conduct experimental evaluations on five representative public datasets with diverse neuronal structures, imaging modalities, and spatial resolutions. Due to the variation in anatomical scale and voxel size across datasets, we do not enforce a fixed sub-volume size. Instead, we adopt dataset-specific configurations to balance spatial context and computational efficiency.

#### AC3/AC4^1^

This dataset consists of two densely labeled subsets of the Kasthuri dataset (Kasthuri et al., 2015), which was imaged from the mouse somatosensory cortex by scanning EM. AC3 and AC4 consist of 256 and 100 sequential images of size 1024×1024, respectively, at 6 × 6 × 29 *nm*^3^ voxel resolution. To ensure the reliability of the experimental results, we carefully divide them into two parts: the bottom 100 layers of AC3 and the complete 100 layers of AC4 are used to simulate unlabeled data *D*_*u*_; the top 100 layers of AC3 are used for testing. The size of the selected sub-volumes (*D*_*l*_) is set as 18 × 380 × 380. Notably, the myelin regions in AC3/AC4 annotations are ignored, leading to confusion in boundary learning. Therefore, we adopt a revised version of annotation for AC4 with myelin annotation (i.e., the training set of SNEMI^2^) and restrict all algorithms to select sub-volumes (*D*_*l*_) from AC4 only.

#### CREMI ^3^

The CREMI dataset comes from an adult Drosophila brain, imaged using serial section transmission EM at a resolution of 4 × 4 × 40 *nm*^3^. The CREMI dataset comprises three labeled subsets (CREMI-A/B/C), each containing 125 sequential images associated with different regions. Since CREMI-A features a homogenous neuron pattern, our experiments employ the CREMI-B/C dataset. Similarly, we divide them into two parts: the top 60 layers of CREMI-B and CREMI-C are considered unlabeled data, denoted as *D*_*u*_; the bottom 60 layers are used for testing. We select and label sub-volumes (*D*_*l*_) of size 18 × 340 × 340 from *D*_*u*_. Moreover, the unlabeled set of CREMI, i.e., CREMI-B+/C+, is used for the investigation of distribution mismatch in Section 5.1.3.

#### AxonEM-H^4^

AxonEM-H dataset is imaged using automated tape-collecting ultra-microtome scanning EM from layer 2 in the temporal lobe of an adult human, containing nine 50 ×512 ×512 sub-volumes at 8 ×8 ×30 *nm*^3^ resolution. The first six sub-volumes are used as unlabeled data (*D*_*u*_), from which labeled sub-volumes (*D*_*l*_) of size 18 × 340 × 340 are selected, while the remaining ones are reserved for evaluation.

#### Hemi-brain^5^

Hemi-brain is a focused ion beam scanning EM volume of the Drosophila melanogaster central brain, acquired at an isotropic resolution of 8 *nm*. For subsequent experiments, we utilize 6 densely labeled volumes provided in the training set, each with a size of 520 × 520 × 520. We partition these volumes into two groups based on their location in the brain. One is designated as unlabeled data (*D*_*u*_), and the other serves as the test data. We select labeled sub-volumes (*D*_*l*_) sized 18 × 300 × 300 from *D*_*u*_.

Kasthuri^6^. The size of the Kasthuri dataset is 1850 × 10747 × 12895, with a voxel resolution of 6 × 6 × 29 *nm*^3^. Our method is evaluated on a sparsely annotated subset cropped from the Kasthuri15 dataset, measuring 300 ×4096 ×4096. The bounding box of this volume ranges from (1050, 6500, 3200) to (1350, 10596, 7296). In the experiments, the top 200 layers serve as unlabeled data (*D*_*u*_), while the bottom 100 layers are designated as the test data. Since the myelin regions are also ignored in official annotations, we select labeled sub-volumes (*D*_*l*_) sized 18 × 380 × 380 from the training set of SNEMI based on embeddings of the unlabeled volume *D*_*u*_.

### 4.2 Evaluation Metrics

Two widely-used metrics for neuron segmentation (Funke et al., 2018; Sheridan et al., 2022; Lee et al., 2021; Huang et al., 2022), the variation of information VI (Nunez-Iglesias et al., 2013) and adapted Rand error ARAND (Arganda-Carreras et al., 2015), are adopted to evaluate the results. VI is defined as a sum of the conditional entropies H(· | ·) between the segmentation proposal A and the ground truth *T*, and can be broken down into two components as

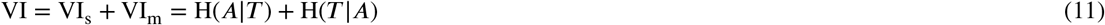

where VI_*s*_ and VI_*m*_, corresponding to split and merge errors, respectively. Define *p*_*ij*_ as the probability that a randomly selected voxel belongs to the segment *i* in *A* and the segment *j* in *T*, the ARAND is formally given by

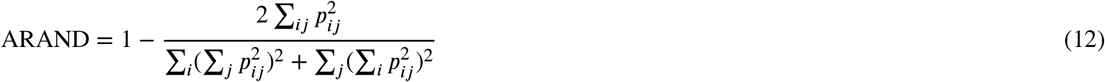

Lower values indicate higher segmentation quality in both metrics.

### 4.3 Implementation Details

All models are trained on 2 NVIDIA RTX 3090 GPUs using Adam optimizer. The input patch size is fixed at 18 × 160 × 160, the batch size is 4, and the learning rate is set as 1e-4 during unsupervised and semi-supervised training. For selective labeling, the unlabeled volume is partitioned using a sliding window approach with a stride of 8 × 40 × 40. The dimension of the embedding space *L* is 80, and the number of neighbors *K* in CCR is set as 30. For semi-supervised learning, the segmentation network is the residual symmetric U-Net (RSU-Net) proposed in (Lee et al., 2017) that is widely adopted in neuron segmentation. We employ the rectangle and quarter masking strategies in axial-through spatial mixing. The weighting coefficient λ for the unlabeled loss component is set to 0.2. The affinity graph has three channels (i.e., *C* = 3), each representing the likelihood between a pixel and its nearest neighbors along the *z, x*, and *y* directions. Once the affinity map is predicted, we obtain the instance segmentation with two commonly adopted approaches, Multicut (Beier et al., 2017) and Waterz (Funke et al., 2018).

## 5. Results

In this section, we first verify the effectiveness of CGS and IIC-Net separately, and then explore their cooperation for more effective semi-supervised segmentation. Finally, we validate the improved pipeline in a large-scale scenario.

### 5.1 Effectiveness of Coverage-based Selective Labeling for Sub-volumes

#### 5.1.1 Comparison with State-of-the-Art Methods

We quantitatively evaluate all strategies, including patch-level and sub-volume selection methods, by training supervised models on the selected data and comparing their segmentation performance. To ensure a fair comparison, all baseline methods are evaluated under an identical labeling budget for each dataset.

For patch-level selective labeling, we consider the following competitors: (a) Random sampling, we repeat the selection process four times with different random seeds and report the average results, denoted as Random_p_. (b) Equispaced sampling is denoted as Equispaced_p_. (c) One-shot active learning methods include *K*-median (Kolluru et al., 2021), *K*-means (Arthur and Vassilvitskii, 2007), farthest point sampling (FPS) (Moenning and Dodgson, 2003; Jin et al., 2022), representative annotation (RA) (Zheng et al., 2019), and unsupervised selective labeling (USL) (Wang et al., 2022a). For sub-volume selection methods, we only compare CGS with random (Random_v_) and equispaced (Equispaced_v_) selection due to the lack of related research. Similarly, we repeat the random selection process four times and calculate the average value. Note that we also compare with CGS’s selection under the fixed-distance coverage function (i.e., each patch can represent patches within a fixed radius in the embedding space), denoted as CGS (FD).

Quantitative results in Table 1, along with the qualitative visualizations in Figure 4, jointly support the following conclusions: (1) As a greedy yet efficient heuristic, CGS enables linear-time sub-volume selection while consistently capturing the underlying data distribution, demonstrating superior performance and strong adaptability across diverse datasets. (2) Sub-volume annotation substantially outperforms patch-level selection, underscoring its superiority for 3D neuron segmentation. Remarkably, even randomly sampled sub-volumes surpass most patch-level selection baselines, revealing the inherent limitations of conventional OSAL methods when directly applied to volumetric EM data. (3) The fixed-distance coverage variant, CGS (FD), overlooks the variable density of patch embeddings in feature space, resulting in suboptimal coverage and degraded performance compared to the *K*-nearest neighbor strategy.

**Table 1.** The quantitative comparisons of selective labeling strategies on the AC3/AC4 and CREMI datasets.

| Methods |  |  | Multicut Post-processing |  |  |  | Waterz Post-processing |  |  |  |
| --- | --- | --- | --- | --- | --- | --- | --- | --- | --- | --- |
|  |  |  | VI<sub>s</sub> ↓ | VI<sub>m</sub> ↓ | VI ↓ | ARAND ↓ | VI<sub>s</sub> ↓ | VI<sub>m</sub> ↓ | VI ↓ | ARAND ↓ |
| AC3/AC4 | Patch-level | Random<sub>p</sub> | 0.9042 | 0.3883 | 1.2925 | 0.1477 | 0.9822 | 0.3837 | 1.3659 | 0.1637 |
|  |  | Equispaced<sub>p</sub> | 0.9459 | 0.3270 | 1.2729 | 0.1224 | 0.9441 | 0.3249 | 1.2690 | 0.1220 |
|  |  | *K*-means | 0.9818 | 0.5333 | 1.5152 | 0.2147 | 0.9859 | 0.4220 | 1.4079 | 0.1401 |
|  |  | *K*-media | 0.9826 | 0.3037 | 1.2863 | 0.1245 | 0.9467 | 0.3692 | 1.3159 | 0.1602 |
|  |  | FPS | 0.9594 | 0.3583 | 1.3177 | 0.1225 | 0.9314 | 0.3769 | 1.3083 | 0.1177 |
|  |  | RA | 0.9424 | 0.3091 | 1.2515 | 0.1158 | 0.9212 | 0.2999 | 1.2211 | 0.1193 |
|  |  | USL | 0.8997 | 0.3507 | 1.2504 | 0.1201 | 0.8974 | 0.3678 | 1.2651 | 0.1288 |
|  | Sub-volume | Random<sub>v</sub> | 0.8774 | 0.3256 | 1.2030 | 0.1182 | 0.8915 | 0.3417 | 1.2307 | 0.1432 |
|  |  | Equispaced<sub>v</sub> | 0.8407 | 0.3430 | 1.1837 | 0.1189 | 0.8520 | 0.3986 | 1.2506 | 0.1499 |
|  |  | CGS (FD) | 0.8733 | 0.2888 | 1.1622 | 0.1150 | 0.9000 | 0.2958 | 1.1958 | 0.1239 |
|  |  | CGS (ours) | 0.8546 | 0.2717 | **1.1263** | **0.1103** | 0.7705 | 0.3196 | **1.0901** | **0.1144** |
|  |  | Improvement (%) | - | - | +6.4% | +6.7% | - | - | +11.4% | +20.1% |
| CREMI | Patch-level | Random<sub>p</sub> | 1.0034 | 0.9599 | 1.9633 | 0.2280 | 1.2531 | 1.4793 | 2.7324 | 0.3742 |
|  |  | Equispaced<sub>p</sub> | 1.0014 | 0.8144 | 1.8158 | 0.2026 | 1.0643 | 1.1739 | 2.2382 | 0.3161 |
|  |  | *K*-means | 1.0656 | 0.8106 | 1.8761 | 0.2240 | 1.1551 | 1.2520 | 2.4070 | 0.3061 |
|  |  | *K*-media | 0.9511 | 0.7897 | 1.7407 | 0.2034 | 1.1473 | 1.3617 | 2.5090 | 0.3919 |
|  |  | FPS | 1.0391 | 0.8372 | 1.8763 | 0.1817 | 1.2892 | 1.3643 | 2.6706 | 0.3406 |
|  |  | RA | 0.9810 | 0.7769 | 1.7579 | 0.1790 | 1.2552 | 1.5709 | 2.8261 | 0.4077 |
|  |  | USL | 1.0014 | 0.8144 | 1.8158 | 0.2026 | 1.0643 | 1.1739 | 2.2381 | 0.3161 |
|  | Sub-volume | Random<sub>v</sub> | 0.9775 | 0.7039 | 1.6814 | 0.1843 | 1.0961 | 1.2460 | 2.3421 | 0.3360 |
|  |  | Equispaced<sub>v</sub> | 0.8838 | 0.7897 | 1.6735 | 0.1626 | 1.0758 | 1.0967 | 2.1725 | 0.3219 |
|  |  | CGS (FD) | 0.9463 | 0.5881 | 1.5326 | 0.1503 | 1.0152 | 1.0728 | 2.0880 | 0.2595 |
|  |  | CGS (ours) | 0.9169 | 0.5675 | **1.4843** | **0.1281** | 0.9693 | 0.8520 | **1.8208** | **0.1944** |
|  |  | Improvement (%) | - | - | +11.7% | +30.5% | - | - | +22.3% | +14.2% |

**Bold** and underlined items indicate the first and second highest scoring results, respectively. We also demonstrate the performance improvements of CGS (ours) over random sub-volume selection.

**Figure 4.**
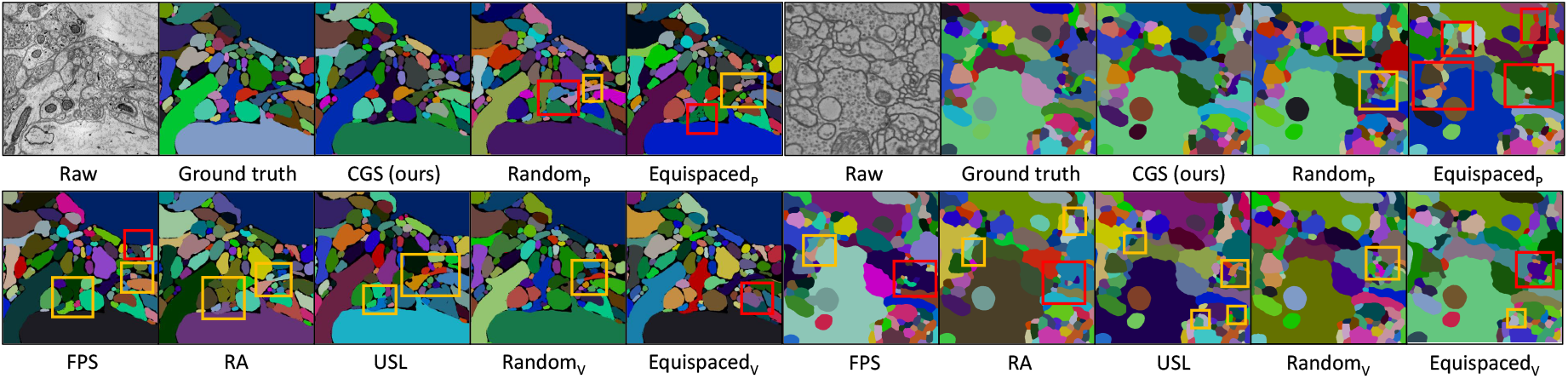
Visual comparison of selective labeling methods on AC3/AC4 (left) and CREMI (right). Red and yellow boxes represent merge and split errors, respectively.

We further investigate the impact of sub-volume size on segmentation performance by comparing CGS with andom_v_. To ensure statistical reliability, we report the average results over four random trials. The quantitative results in TABLE 2 reveal several important insights. First, CGS consistently outperforms random selection across all sub-volume sizes, demonstrating both effectiveness and robustness. Second, sub-volume size appears to influence labeling efficiency, particularly under random selection. Smaller sizes may struggle to capture the morphological continuity of neurons, while larger sizes can limit the number of candidate regions, potentially reducing semantic coverage.

**Table 2.** Quantitative Comparisons on AC3/AC4 dataset with different sub-volume sizes with the same labeling budget.

| Size | Methods | Multicut |  | Waterz |  |
| --- | --- | --- | --- | --- | --- |
|  |  | VI ↓ | ARAND ↓ | VI ↓ | ARAND ↓ |
| *Patch-level Annotation* |  |  |  |  |  |
| 18×160×160 | Random<sub>p</sub> | 1.2925 | 0.1477 | 1.3659 | 0.1637 |
| *Sub-volume Annotation* |  |  |  |  |  |
| 18×220×220 | Random<sub>v</sub> | 1.2254 | 0.1432 | 1.2838 | 0.1538 |
|  | CGS | **1.1524** | **0.1138** | **1.1870** | **0.1170** |
| 18×300×300 | Random<sub>v</sub> | 1.1958 | 0.1214 | 1.2127 | 0.1406 |
|  | CGS | **1.1484** | **0.1111** | **1.1241** | **0.1121** |
| 18×380×380 | Random<sub>v</sub> | 1.2030 | 0.1182 | 1.2307 | 0.1432 |
|  | CGS | **1.1263** | **0.1103** | **1.0901** | **0.1144** |
| 18×460×460 | Random<sub>v</sub> | 1.1620 | 0.1348 | 1.2445 | 0.1744 |
|  | CGS | **1.1024** | **0.1163** | **1.1210** | **0.1130** |

#### 5.1.2 Visualization of CGS in Feature Space

We employ the UMAP to project the patch embeddings for the AC3/AC4 dataset into a two-dimensional feature space. Clustering those vectors allows for identifying biologically meaningful distinctions. We visualize the selected sub-volumes along with their associated patch embeddings, i.e., the selected and covered vectors. As shown in Figure 5, representative neuronal structures in the raw volume are semantically covered by the selected sub-volumes, including myelin, dendrite shaft (or soma), dendrite spine, mitochondria, vesicle cloud, and axon. This visualization validates the effectiveness of CGS in extracting informative and diverse sub-regions from EM volumes.

**Figure 5.**
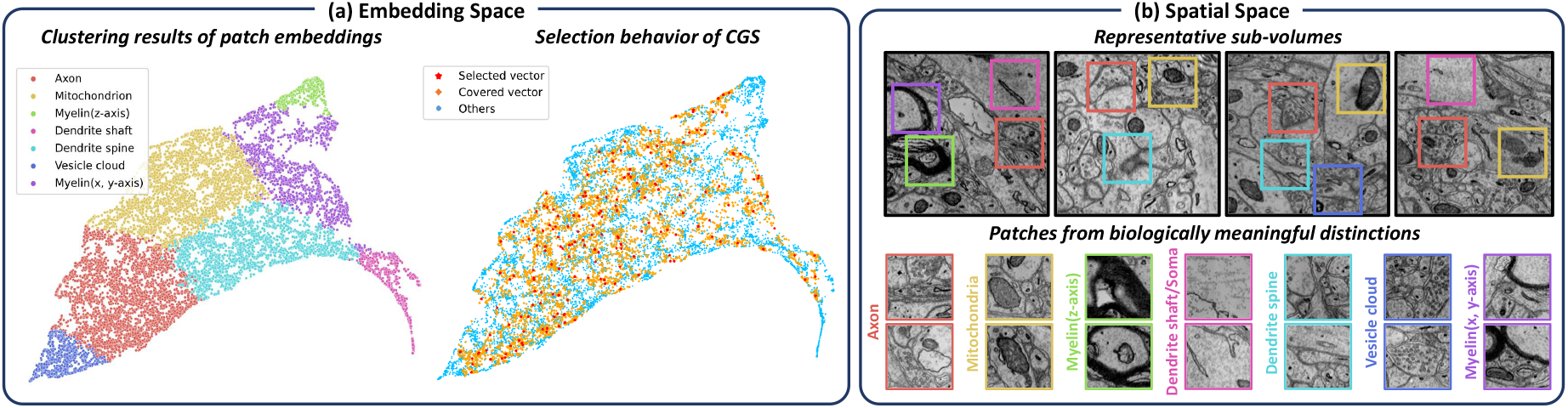
(a) UMAP visualization of patch embeddings from the AC3/AC4 dataset. Spectral clustering reveals biologically meaningful groups, with dominant structures labeled in the legend. CGS-selected patches are well distributed across the embedding space. (b) Spatial view of selected sub-volumes and example patches corresponding to distinct biological structures.

#### 5.1.3 Impact of Data-level Distribution Mismatch on SSL

This subsection investigates the impact of data-level distribution mismatch in semi-supervised neuron segmentation by conducting controlled experiments using the proposed IIC-Net on the AC3/AC4 and CREMI datasets. Building on the conclusions in Table 1 and Figure 5, CGS-selected sub-volumes (*D*_*l*_) are well aligned with the unlabeled data (*D*_*u*_). To introduce distribution mismatch for comparison, we design two variants: (1) Fix *D*_*u*_, and replace *D*_*l*_ with randomly cropped sub-volumes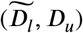, repeated four times under the same labeling budget. (2) Fix *D*_*l*_ and randomly alter the unlabeled set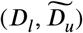. For AC3/AC4, we use eight volumes from the Kasthuri dataset (100×1024×1024); for CREMI, we sample eight sub-volumes from CREMI-B+/C+ (60 slices each). Each setting also includes four randomly sampled subsets to ensure statistical reliability. The quantitative findings in Table 3 highlight the adverse effect of distribution mismatch on semi-supervised neuron segmentation. This subsection fundamentally confirms the core motivation and necessity of our distribution-aware design at the data level.

**Table 3.**
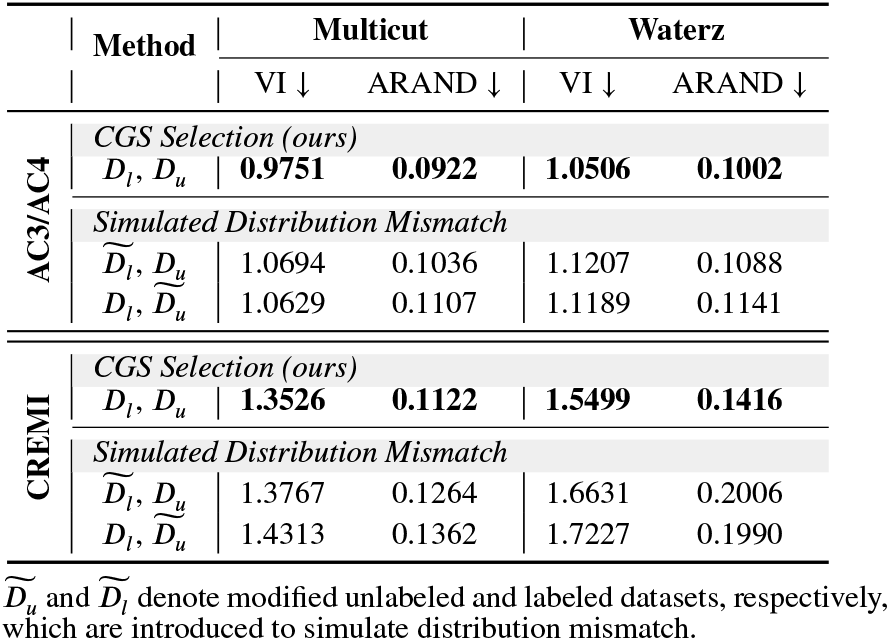
Impact of distribution mismatch in semi-supervised learning on the AC3/AC4 and CREMI datasets.

|  | Method | Multicut |  | Waterz |  |
| --- | --- | --- | --- | --- | --- |
|  |  | VI ↓ | ARAND ↓ | VI ↓ | ARAND ↓ |
| AC3/AC4 | *CGS Selection (ours)* |  |  |  |  |
| | $D_l, D_u$ | **0.9751** | **0.0922** | **1.0506** | **0.1002** |
|  | *Simulated Distribution Mismatch* |  |  |  |  |
| | $\tilde{D}_l, D_u$ | 1.0694 | 0.1036 | 1.1207 | 0.1088 |
| | $D_l, \tilde{D}_u$ | 1.0629 | 0.1107 | 1.1189 | 0.1141 |
| CREMI | *CGS Selection (ours)* |  |  |  |  |
| | $D_l, D_u$ | **1.3526** | **0.1122** | **1.5499** | **0.1416** |
|  | *Simulated Distribution Mismatch* |  |  |  |  |
| | $\tilde{D}_l, D_u$ | 1.3767 | 0.1264 | 1.6631 | 0.2006 |
| | $D_l, \tilde{D}_u$ | 1.4313 | 0.1362 | 1.7227 | 0.1990 |

$\tilde{D}_u$ and $\tilde{D}_l$ denote modified unlabeled and labeled datasets, respectively, which are introduced to simulate distribution mismatch.

#### 5.2. Effectiveness of Mixed-view Consistency Learning

#### 5.2.1 Comparison with State-of-the-Art Methods

In this subsection, we verify the state-of-the-art performance of the proposed semi-supervised framework, i.e., IIC-Net. Experiments on AC3/AC4 and CREMI datasets are conducted on CGS-selected sub-volumes *D*_*l*_ and unlabeled data *D*_*u*_. Two supervised learning methods and six semi-supervised learning methods are adopted for comparison, including (1) RSU-Net (Lee et al., 2017), using only labeled data in a supervised manner, (2) RSU-Net w/ Mixing, incorporating axial-through spatial mixing for labeled data, (3) PseudoSeg (Zou et al., 2020), a classical semi-supervised approach that enforces consistency between unlabeled data and hard pseudo labels, (4) UA-MT (Yu et al., 2019), utilizing plausible predictions to guide consistency learning, (5) SASSNet (Li et al., 2020), employs an adversarial learning strategy to enforce a geometric shape constraint between the affinity maps of labeled and unlabeled data, (6) MC-Net (Wu et al., 2021), proposes mutual consistency regulation to encourage low-entropy predictions, (7) BCP (Bai et al., 2023), propose bidirectional copy-paste method within a Mean Teacher framework, and (8) SSNS-Net (Huang et al., 2022), the first semi-supervised neuron segmentation framework introducing innovative network initialization and modeling consistency between soft pseudo labels and perturbed predictions of unlabeled data. Additionally, we perform an ablation study to demonstrate the benefits of intra-and inter-dataset mixing, respectively.

We consider the following variants: IIC-Net w/o Inter-mixing, where we disable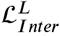and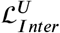, and IIC-Net w/o Intra-mixing, where Π in M_1_ and M_3_ are set as **1**. For a fair comparison, we use consistent network architecture and augmentation strategies across all baseline methods.

The quantitative results under two post-processing algorithms present in Table 4 support the following conclusions:(1) Performing spatial mixing on labeled data (RSU-Net w/ Mix) can directly improve the segmentation performance in the supervised setting. This reveals the effectiveness of mixing strategies in promoting a better understanding of neuron boundaries. (2) The proposed IIC-Net achieves consistent improvements over all baselines, which confirms the effectiveness of mixed-view consistency regularization. (3) The performance of IIC-Net decreases when intra-dataset mixing is turned off (w/o Intra-mix), emphasizing its critical role in facilitating adaptation to mixed inputs. Notably, when inter-dataset mixing (w/o Inter-mix) is disabled, the performance drops even more. Such results suggest that inter-mixing serves as a key factor. It connects labeled and unlabeled datasets and encourages the network to learn shared semantics. We also provide a visual comparison of different semi-supervised methods in Figure 6.

**Table 4.** The quantitative comparisons of semi-supervised methods on the AC3/AC4 and CREMI datasets.

| Methods |  |  | Multicut Pose-processing |  |  |  | Waterz Pose-processing |  |  |  |
| --- | --- | --- | --- | --- | --- | --- | --- | --- | --- | --- |
|  |  |  | VI<sub>s</sub> ↓ | VI<sub>m</sub> ↓ | VI ↓ | ARAND ↓ | VI<sub>s</sub> ↓ | VI<sub>m</sub> ↓ | VI ↓ | ARAND ↓ |
| AC3/AC4 | Sup. | RSU-Net | 0.8546 | 0.2717 | 1.1263 | 0.1103 | 0.7705 | 0.3196 | 1.0901 | 0.1144 |
|  |  | RSU-Net w/ Mix | 0.7877 | 0.3482 | 1.1059 | 0.1061 | 0.7983 | 0.3144 | 1.1127 | 0.1031 |
|  | Semi. | PseudoSeg | 0.8190 | 0.2687 | 1.0876 | 0.1073 | 0.8489 | 0.2934 | 1.1424 | 0.1270 |
|  |  | UA-MT | 0.8018 | 0.2645 | 1.0662 | 0.0998 | 0.8588 | 0.2886 | 1.1473 | 0.1172 |
|  |  | SASSNet | 0.7784 | 0.2914 | 1.0698 | 0.1143 | 0.8434 | 0.2467 | 1.0902 | 0.1086 |
|  |  | MC-Net | 0.7945 | 0.2592 | 1.0538 | 0.1082 | 0.8329 | 0.2390 | 1.0719 | 0.1049 |
|  |  | BCP | 0.7864 | 0.2771 | 1.0634 | 0.1030 | 0.8202 | 0.3035 | 1.1238 | 0.1104 |
|  |  | SSNS-Net | 0.7308 | 0.3166 | 1.0473 | 0.1025 | 0.7781 | 0.3113 | 1.0894 | 0.1165 |
|  |  | IIC-Net w/o Inter-mix | 0.7395 | 0.2899 | 1.0293 | 0.1008 | 0.7732 | 0.2882 | 1.0614 | **0.0960** |
|  |  | IIC-Net w/o Intra-mix | 0.7335 | 0.2792 | 1.0127 | 0.0937 | 0.8136 | 0.2559 | 1.0695 | 0.1072 |
|  |  | IIC-Net (ours) | 0.6895 | 0.2855 | **0.9751** | **0.0922** | 0.7813 | 0.2693 | **1.0506** | <u>0.1002</u> |
|  | Improvement (%) |  | - | - | +13.4% | +16.4% | - | - | +3.6% | +12.4% |
| CREMI | Sup. | RSU-Net | 0.9169 | 0.5675 | 1.4843 | 0.1281 | 0.9693 | 0.8520 | 1.8208 | 0.1944 |
|  |  | RSU-Net w/ Mix | 0.8462 | 0.6004 | 1.4466 | 0.1336 | 0.9451 | 0.8743 | 1.8193 | 0.2034 |
|  | Semi. | PseudoSeg | 0.9068 | 0.5406 | 1.4474 | 0.1362 | 0.9819 | 0.8659 | 1.8478 | 0.2289 |
|  |  | UA-MT | 0.8966 | 0.5420 | 1.4386 | 0.1216 | 0.9718 | 0.7593 | 1.7311 | 0.1730 |
|  |  | SASSNet | 0.8917 | 0.5552 | 1.4469 | 0.1265 | 0.9804 | 0.7711 | 1.7514 | 0.1815 |
|  |  | MC-Net | 0.8970 | 0.5409 | 1.4379 | 0.1235 | 0.9678 | 0.7537 | 1.7215 | 0.1942 |
|  |  | BCP | 0.9134 | 0.5229 | 1.4363 | 0.1392 | 0.9279 | 0.7063 | 1.6342 | 0.1697 |
|  |  | SSNS-Net | 0.8922 | 0.5329 | 1.4251 | 0.1217 | 0.9451 | 0.7229 | 1.6679 | 0.1633 |
|  |  | IIC-Net w/o Inter-mix | 0.8472 | 0.5367 | 1.3838 | 0.1229 | 0.9323 | 0.7099 | 1.6400 | 0.1831 |
|  |  | IIC-Net w/o Intra-mix | 0.8767 | 0.4986 | 1.3753 | 0.1258 | 0.8886 | 0.6707 | 1.5593 | 0.1576 |
|  |  | IIC-Net (ours) | 0.8534 | 0.4992 | **1.3526** | **0.1122** | 0.8669 | 0.6830 | **1.5499** | **0.1416** |
|  | Improvement (%) |  | - | - | +8.9% | +12.4% | - | - | +14.9% | +27.2% |

Sup. and Semi. are abbreviations for supervised and semi-supervised learning. **Bold** and underlined items indicate the first and second scoring results, respectively. We further quantify the performance improvements of IIC-Net (ours) compared to the supervised baseline (RSU-Net).

**Figure 6.**
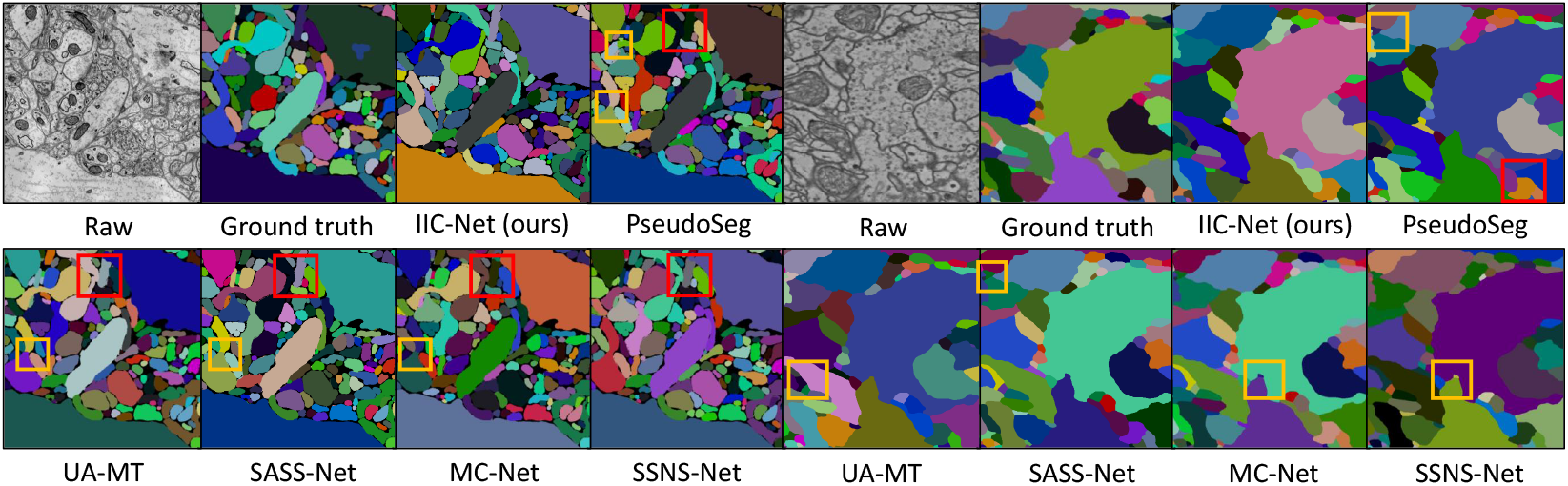
Qualitative comparison of semi-supervised methods on the AC3/AC4 (left) and CREMI (right) dataset. The red and yellow boxes represent merge and split errors, respectively.

#### 5.2.2 Visualization of IIC-Net in Feature Space

To further assess the effectiveness of IIC-Net at the model level, we visualize the deep feature distributions of labeled and unlabeled data using kernel density estimation under different training methods, and quantify their similarity via Jensen–Shannon distance. Figure 7 demonstrates that semi-supervised models achieve significantly enhanced congruence between the feature distributions of labeled and unlabeled data compared to the supervised baseline. IIC-Net, in particular, demonstrates superior alignment compared to SSNS-Net (Huang et al., 2022), underscoring the effectiveness of mixed-view consistency learning in promoting seamless feature transitions between labeled and unlabeled data, thereby fostering more generalizable semi-supervised learning.

**Figure 7.**
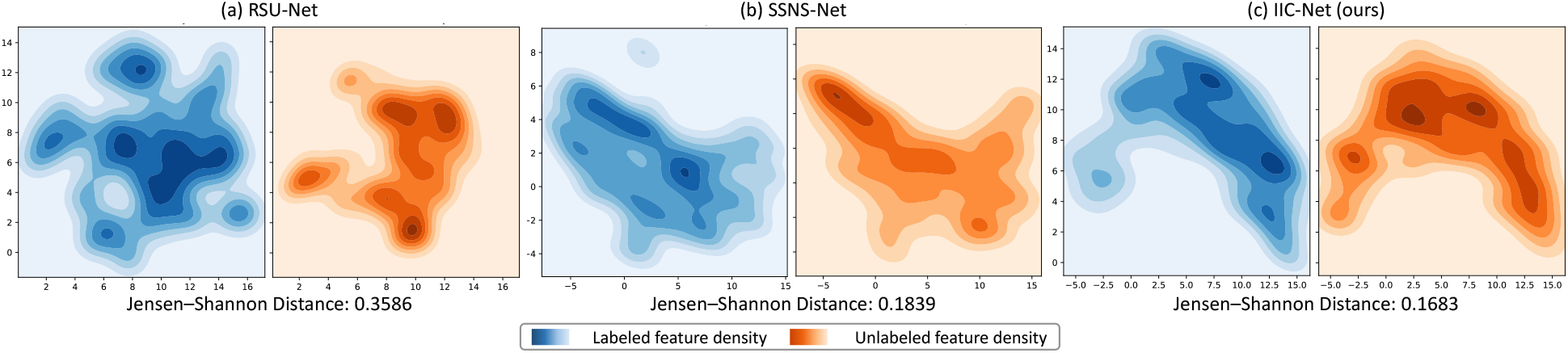
Kernel density estimations of deep features extracted from labeled and unlabeled data across different training methods.

### 5.3 Ablation Study for the Distribution-aware Semi-supervised Pipeline

To evaluate the effectiveness of our pipeline, we perform an ablation study analyzing the interplay between selective labeling and semi-supervised training for neuron segmentation. Experiments are conducted on three datasets from different species: rat (AC3/AC4), fly (Hemi-brain), and human (AxonEM-H). We progressively increase labeling costs and assess performance under both supervised and semi-supervised settings.

For selective labeling, we compare CGS against random sub-volume selection, reporting the best result across four runs, denoted as Random^*^. For semi-supervised learning, we adopt SSNS-Net (Huang et al., 2022), a representative baseline, for comparison. Results using Multicut post-processing are shown in Figure 8. Our framework, which combines CGS-based sub-volume selection with IIC-Net training, consistently outperforms all baselines across various annotation budgets and datasets. Notably, our method achieves performance comparable to full supervision using only 10% labeled data on the AC3/AC4 dataset. Moreover, 3D visualizations in Figure 9 illustrate the pipeline’s superior ability to preserve fine-grained neuronal connectivity.

**Figure 8.**
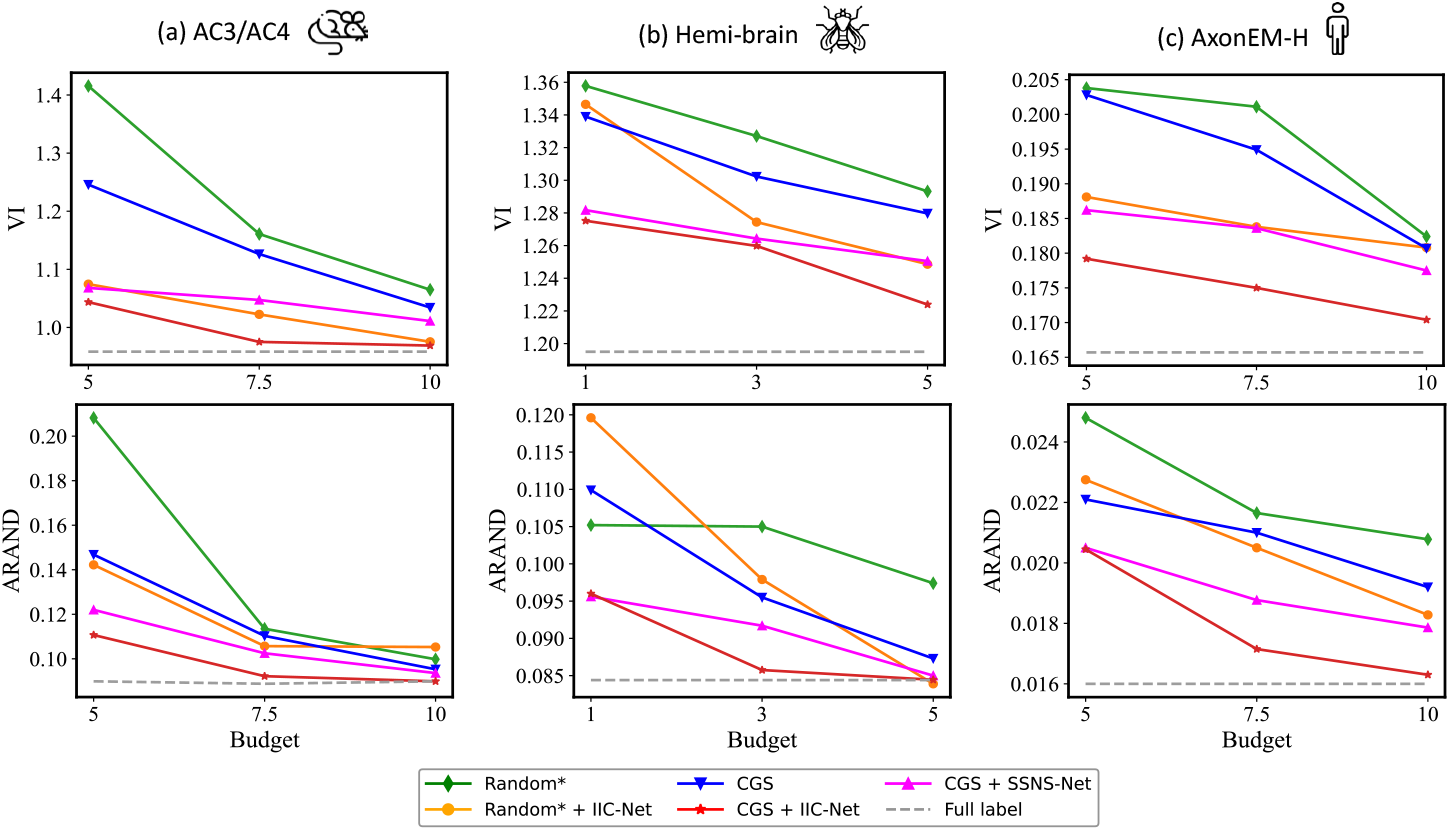
Quantitative Comparisons on the AC3/AC4, Hemi-brain and AxonEM-H dataset with different annotation budgets.

**Figure 9.**
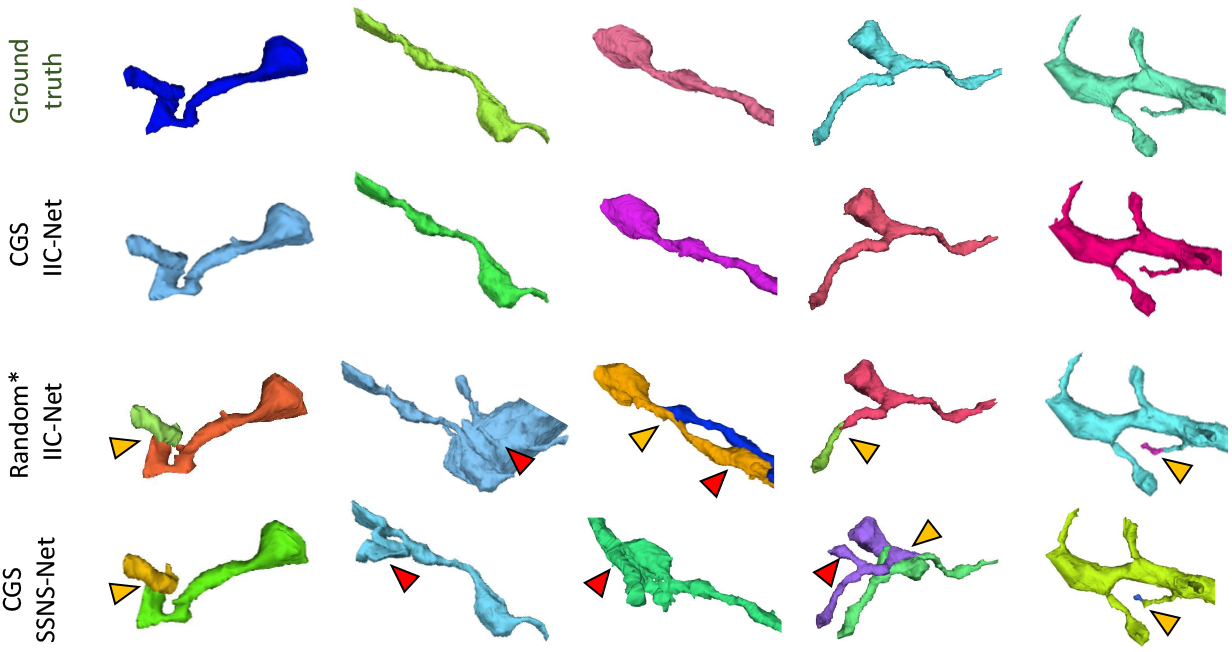
3D visual comparison of segmentation results on the AC3/AC4 dataset with labeling cost of 7.5%. The red and yellow arrows represent merge and split errors, respectively.

### 5.4 Evaluation on Large-scale EM Data

We apply the proposed pipeline—selective labeling via CGS combined with semi-supervised training via IIC-Net—on the Kasthuri dataset to evaluate its practical utility in large-scale neuron reconstruction. Quantitative results are shown in TABLE 5. Quantitative results are summarized in TABLE 5. Given the same labeling budget, including the full 100%, IIC-Net consistently outperforms the supervised baseline in segmentation accuracy. Considering the significant time and cost of expert proofreading, semi-supervised training offers more reliable predictions without additional labeling or inference overhead, making it highly promising for large-scale connectomics. Notably, the fully supervised approach (VI: 0.7751, ARAND: 0.1731) is effectively matched by our pipeline using only 20% labeled data (VI: 0.7842, ARAND: 0.1710), demonstrating its effectiveness for cost-effective neuron segmentation.

**Table 5.** Evaluation on Kasthuri15 using the proposed semi-supervised pipeline.

| Budget | SL | SSL | VI ↓ | ARAND ↓ |
| --- | --- | --- | --- | --- |
| 10% | CGS | ✗ | 0.8540 | 0.1810 |
|  | CGS | IIC-Net | **0.8100** | **0.1741** |
|  | Improvement (%) |  | +5.15% | +3.81% |
| 20% | CGS | ✗ | 0.8236 | 0.1795 |
|  | CGS | IIC-Net | **0.7842** | **0.1710** |
|  | Improvement (%) |  | +4.78% | +4.74% |
| 100% | ✗ | ✗ | 0.7751 | 0.1731 |
|  | ✗ | IIC-Net | **0.7362** | **0.1632** |
|  | Improvement (%) |  | +5.02% | +5.72% |

SL and SSL denote selective labeling and semi-supervised learning, respectively.

## 6 Conclusion

Enhancing segmentation accuracy at the automated stage is critical for scalable and cost-effective connectomics. Semi-supervised learning offers a compelling strategy by exploiting abundant unlabeled data without incurring additional annotation or inference overhead. However, a fundamental challenge remains: distribution mismatch between the limited labeled subset and the broader, heterogeneous unlabeled data severely hampers model generalization. To address this, we propose a novel semi-supervised pipeline for 3D neuron segmentation in EM volumes. Our approach combines coverage-guided selective labeling with mixed-view consistency regularization, effectively aligning labeled and unlabeled distributions at both the data and model levels. Extensive experiments across varying species and resolutions demonstrate the superior performance and generalizability of the proposed framework, underscoring its potential to reduce manual effort and accelerate large-scale connectomic analyses.

## Acknowledgments

This work was supported in part by the STI 2030–Major Projects under Grant 2021ZD0204500 and Grant 2021ZD0204503, in part by the National Natural Science Foundation of China under Grant 32171461, and in part by the Beijing Natural Science Foundation under Grant 5254042.

## Footnotes

1 https://lichtman.rc.fas.harvard.edu/vast/AC3AC4Package.zip

2 ^2^https://snemi3d.grand-challenge.org

3 ^3^https://cremi.org

4 ^4^https://axonem.grand-challenge.org

5 ^5^https://github.com/funkelab/lsd#notebooks

6 ^6^https://neurodata.io/data/kasthuri15

